# Carbapenem-Resistant *Acinetobacter baumannii* in US hospitals: diversification of circulating lineages and antimicrobial resistance

**DOI:** 10.1101/2021.09.21.461323

**Authors:** Alina Iovleva, Mustapha M. Mustapha, Marissa P. Griffith, Lauren Komarow, Courtney Luterbach, Daniel R Evans, Eric Cober, Sandra S. Richter, Kirsten Rydell, Cesar A. Arias, Jesse T. Jacob, Robert A. Salata, Michael J. Satlin, Darren Wong, Robert A. Bonomo, David van Duin, Vaughn S. Cooper, Daria Van Tyne, Yohei Doi

## Abstract

Carbapenem-resistant *Acinetobacter baumannii* (CR*Ab*) are a major cause of healthcare-associated infections. CR*Ab* are typically multidrug-resistant and infection is difficult to treat. Despite the urgent threat that CR*Ab* pose, few systematic studies of CR*Ab* clinical and molecular epidemiology have been conducted. The Study Network of *Acinetobacter* as a Carbapenem-Resistant Pathogen (SNAP) is designed to investigate the clinical characteristics and contemporary population structure of CR*Ab* circulating in US hospital systems using whole genome sequencing (WGS). Analysis of the initial 120 SNAP patients from four US centers revealed that CR*Ab* remain a significant threat to hospitalized patients, affecting the most vulnerable patients and resulting in 24% all-cause 30-day mortality. The majority of currently circulating isolates belonged to ST2^Pas^, a part of Clonal Complex 2 (CC2), which is the dominant drug-resistant lineage in the United States and Europe. We identified three distinct sub-lineages within CC2, which differed in their antibiotic resistance phenotypes and geographic distribution. Most concerning, colistin resistance (38%) and cefiderocol (10%) resistance were common within CC2 sub-lineage C (CC2C), where the majority of isolates belonged to ST2^Pas^/ST281^Ox^. Additionally, we identified a newly emergent lineage, ST499^Pas^ that was the most common non-CC2 lineage in our study and had a more favorable drug susceptibility profile compared to CC2. Our findings suggest a shift within the CR*Ab* population in the US during the past 10 years, and emphasize the importance of real-time surveillance and molecular epidemiology in studying CR*Ab* dissemination and clinical impact.

**Importance:** Carbapenem-resistant *Acinetobacter baumannii* (CR*Ab*) constitute a major threat to public health. To elucidate the molecular and clinical epidemiology of CR*Ab* in the US, clinical CR*Ab* isolates were collected along with data on patient characteristics and outcomes and bacterial isolates underwent whole genome sequencing and antibiotic susceptibility phenotyping. Key findings included emergence of new sub-lineages within the globally predominant clonal complex (CC) 2, increased colistin and cefiderocol resistance within one of the CC2 sub-lineages, and the emergence of ST499^Pas^ as a previously unrecognized CR*Ab* lineage in US hospitals.

## Introduction

Carbapenem-resistant *Acinetobacter baumannii* (CR*Ab*) constitute a major threat to public health. CR*Ab* are extensively resistant to multiple antimicrobial agents, often spread among hospitalized patients, and cause difficult-to-treat infections associated with high mortality (1). The World Health Organization (WHO) and the Centers for Disease Control and Prevention (CDC) have designated CR*Ab* a “priority pathogen” based on the lack of effective treatment options, and have pointed to an urgent need for additional research (2-4). Prior studies have shown that several genetically distinct clonal *A. baumannii* lineages/groups are currently circulating around the world, with the three most prevalent global lineages referred to as Clonal Complex (CC) 1, CC2, and CC3. These designations are reflected in the multi-locus sequence types of each lineage (ST1, ST2, and ST3, respectively) as defined by the commonly used Pasteur Institute scheme. ST2^Pas^ is the most common CC2 lineage in the US, and other CC2 and non-CC2 lineages are found less commonly (5, 6). Our previous analysis at a single health system in Pennsylvania found substantial genetic diversity within extensively drug-resistant ST2^Pas^ *A. baumannii*, which could be grouped into multiple distinct sub-lineages by MLST and WGS analysis (7). Despite being a major public health concern, our current understanding of the CR*Ab* lineages and sub-lineages circulating in the United States is limited. Systematic studies of CR*Ab* are of paramount importance in devising strategies to prevent their dissemination and improve clinical outcomes.

The Study Network of *Acinetobacter* as a Carbapenem-Resistant Pathogen (SNAP) is a prospective, observational, multicenter clinical study that is designed to elucidate the clinical characteristics, treatment outcomes, and contemporary genomic epidemiology of CR*Ab* through consecutive enrollment of hospitalized patients with clinical cultures positive for CR*Ab* at multiple health systems throughout the US. In this analysis, we describe the results from this effort, comprising 120 unique patients and 150 CR*Ab* isolates collected during the first year of the study from four health systems in the US, with a focus on patient characteristics, bacterial population structure and antibiotic resistance profiles.

## Methods

### Patients

Patients were included in the study if CR*Ab* were isolated in a clinical culture from any anatomic site during hospitalization between September 2017 and October 2018. Carbapenem resistance was determined based on the Clinical and Laboratory Standards Institute (CLSI) interpretive criteria for meropenem or imipenem non-susceptibility (minimum inhibitory concentration [MIC], ≥4 mg/L). A total of twenty-three hospitals in four quaternary health systems (Cleveland Clinic Foundation, University of Pittsburgh Medical Center, University of Texas Health Science Center at Houston, and University of North Carolina Chapel Hill) enrolled patients in study phase. The study was approved by the Institutional Review Boards (IRB) of all the health systems with a waiver of patient consent.

### Clinical information

Clinical data based on electronic health record collected included patient demographics, underlying comorbidities (Charlson comorbidity index [CCI]), the severity of illness as defined by the Pitt bacteremia score, microbiology reports, resolution of infection symptoms, duration of hospital stay, disposition after discharge, readmission at 90 days, mortality at 30 and 90 days, and infection versus colonization status (8, 9). Infection and colonization were defined by previously described criteria, with the exception of respiratory infections, as patients with CR*Ab* respiratory infections do not necessarily meet the criteria outlined by the American Thoracic Society and the Infectious Diseases Society of America (10-12). Respiratory isolates were considered to be causing an infection if the respiratory diagnosis on the case report form was tracheobronchitis, pneumonia without mechanical ventilation, ventilator-associated pneumonia, or an “other” diagnosis after review by two study investigators. All other cultures, including those missing information needed for the assignment of infection/colonization, were considered to represent colonization. The desirability of outcome ranking (DOOR) analysis was used to assess the following deleterious and adverse events: 1) absence of clinical and symptomatic response or relapse of infection; 2) unsuccessful discharge, which included death, discharge to hospice, hospitalization >30 days and readmission; 3) new-onset renal failure within 30 days after the index culture; and 4) *Clostridioides difficile* infection within 30 days after index culture, as described previously (10).

### Microbiology

Bacterial identification and susceptibility testing were performed by each contributing microbiology laboratory using Biotyper (Bruker, Billerica, MA, USA), MicroScan (Beckman Coulter, Atlanta, GA, USA) or VitekMS, Vitek2, Etest (all bioMérieux, Durham, NC, USA), BD Phoenix, BBL disks (both BD, Durham, NC, USA), Sensititre (Thermo Fisher, Waltham, MA, USA), disk diffusion or in-house agar dilution.

At the central research laboratory, MICs of each agent active against *A. baumannii* (amikacin, gentamicin, tobramycin, doxycycline, minocycline, tigecycline, ciprofloxacin, levofloxacin, trimethoprim-sulfamethoxazole, imipenem, meropenem, doripenem, cefepime, ceftazidime, ampicillin-sulbactam, and colistin) were determined using Sensititre™ GNX3F plates (Thermo Fisher Scientific, Waltham, MA). CLSI breakpoints were used to determine susceptibility.

Cefiderocol susceptibility testing was performed using an iron-depleted, cation-adjusted Mueller-Hinton broth microdilution panel (International Health Management Associates, Schaumburg, IL, USA). Cefiderocol MIC results were interpreted using investigational breakpoints with MIC ≤ 4 μg/ml considered susceptible and MIC >4 μg/ml as non-susceptible. As CLSI breakpoints are not available for tigecycline, we defined susceptibility as MIC ≤2 μg/ml, and non-susceptibility as MIC ≥4 μg/ml, based on previous literature (13).

### Whole genome sequencing and phylogenetic analysis

Genomic DNA was extracted from isolates using a DNeasy Blood & Tissue Kit (Qiagen, Germantown, MD). Whole genome sequencing was performed on a NextSeq 550 instrument (Illumina, San Diego, CA), using 2×150-bp paired-end reads at the Microbial Genome Sequencing Center (Pittsburgh, PA). Additionally, five isolates representing major sub-lineages were sequenced with long-read technology on an Oxford Nanopore MinION device (Oxford Nanopore Technologies, Oxford, United Kingdom). Resulting reads were quality processed through our bioinformatics pipeline. Five isolates were excluded from further molecular and antimicrobial susceptibility analysis as they were identified as bacterial species other than *A. baumannii* (4 isolates) or were carbapenem-susceptible *A. baumannii* (1 isolate). Details of sequencing, bioinformatics, and phylogenetic analyses are available in the Supplementary Materials.

### Nucleotide sequence accession numbers

Raw sequence reads and draft genome assemblies have been deposited in the NCBI database under BioProject PRJNA667103, under accession numbers SAMN16351076 - SAMN16351208.

## RESULTS

### Patients and clinical epidemiology

120 unique patients admitted to four health systems in the US in 2017-2018 were enrolled in the first phase of the SNAP cohort (Table 1). In this cohort, 135 admissions were recorded, and 155 clinical cultures yielded CR*Ab* (1-5 isolates per patient). Clinical data were available from all 120 patients. The enrollments were from Cleveland (55%), Pittsburgh (23%), Houston (20%), and Chapel Hill (3%). Median patient age was 61. 60% were male. Most patients had comorbid conditions, with a median Charlson comorbidity score of 3, and 48% were critically ill (Pitt bacteremia score ≥4) at the time of initial CR*Ab* isolation (8). More than half of patients were admitted from long-term care settings, with 41% admitted from long-term care facilities and 11% from long-term acute care hospitals. All-cause mortality rates at 30 and 90 days from the date of index culture collection were 24% and 27%, respectively. 30- and 90-day mortality in those deemed to have infection was 26%. Readmission within 90 days occurred in 54% of cases. Using DOOR outcomes at 30 days after index culture, 44% were alive without events, 19% were alive with one event, 13% were alive with two or three events.

**Table 1.** Clinical characteristics and outcomes following collection of the first CR*Ab* isolate from each patient.

| Characteristics |  | Total (n120) |
| --- | --- | --- |
| Age at culture | Median (IQR) | 61 (51, 70) |
| Gender | Male | 72 (60%) |
|  | Female | 48 (40%) |
| Race | White | 74 (62%) |
|  | Black | 33 (28%) |
|  | Other | 7 (6%) |
|  | Unknown | 6 (5%) |
| CCI* | Median (IQR) | 3 (1, 4) |
| Pitt bacteremia score** | Median (IQR) | 3 (2, 6) |
| Admitted from | Home | 37 (31%) |
|  | Transfer from other hospital | 21 (18%) |
|  | Long term care | 49 (41%) |
|  | Long term acute care (LTAC) | 13 (11%) |
| Study site | Cleveland Clinic Foundation | 66 (55%) |
|  | University of North Carolina at Chapel Hill | 3 (3%) |
|  | University of Pittsburgh Medical Center | 27 (23%) |
|  | University of Texas Hospitals | 24 (20%) |
| Culture | Respiratory infection | 33 (28%) |
|  | Respiratory colonization | 23 (19%) |
|  | Wound infection | 20 (17%) |
|  | Wound colonization | 23 (19%) |
|  | Blood infection | 9 (8%) |
|  | Urine infection | 3 (3%) |
|  | Urine colonization | 6 (5%) |
|  | Other colonization | 2 (2%) |
|  | Non-wound abdominal infection | 1 (1%) |
| DOOR categories at 30 days *** | Alive without events | 53 (44%) |
|  | Alive with one event | 23 (19%) |
|  | Alive with two or three events | 15 (13%) |
|  | Dead | 29 (24%) |
| DOOR events at 30 days | Dead or discharged to hospice | 24 (20%) |
|  | No clinical response | 41 (34%) |
|  | Renal failure | 8 (7%) |
|  | *C. difficile* | 3 (3%) |
| **Mortality** | 30 days | 29 (24%) |
|  | 90 days | 32 (27%) |
| **Mortality among subjects with infection** | 30 days | 17 (26%) |
|  | 90 days | 17 (26%) |
| **Readmission** | 90 days | 51 (54%) |
| **Readmission among subjects with infection** | 90 days | 25 (38%) |

Data are n (%) or median (IQR).
\* Charlson comorbidity index (CCI) is a chronic comorbidity score with a range from 0 to 37, with higher scores indicating more comorbid conditions present. A patient with a score of 3 could have three level 1 comorbid conditions (e.g., dementia, chronic pulmonary disease, and congestive heart failure), one level 1 (e.g., dementia) and one level 2 comorbid condition (e.g., leukemia), or one level 3 condition (moderate or severe liver disease).
\*\* Pitt bacteremia score is an acute severity of illness score. Higher scores indicate more severe illness. A patient with a score of 3 would have one level 1 marker (e.g., disoriented mental status) and one level 2 marker of acute illness (e.g., hypotension).
\*\*\* DOOR, desirability of outcome ranking. DOOR analysis components are defined in the methods section.

Among the 120 enrolled patients, the respiratory tract was the predominant anatomic source of the first available CR*Ab* isolate (47%), followed by wound (36%) (Table 1). 59% of isolates from respiratory source were associated with infection, while 46% of isolates from wound cultures and 33% from urine cultures met the definition of infection.

### Molecular epidemiology and bacterial population structure

To understand the population structure and distribution of CR*Ab* in the US, phylogenetic analyses and multi-locus sequence type (MLST) identification were performed on the first available isolate from each patient.

A whole genome phylogeny showed a diverse population structure (Figure 1). We defined single nucleotide polymorphism (SNP) thresholds that clustered study isolates into clearly defined clonal lineages and sub-lineages and correlated them with established Pasteur (^Pas^) and Oxford (^Ox^) MLST schemes. A cut-off of 10,000 SNPs differentiated CR*Ab* lineages belonging to different Pasteur ST types and CCs. A cut-off of 2,000 SNPs further defined major sub-lineages within the Pasteur STs which largely correlated with Oxford STs. The 115 available isolates belonged to 10 different Pasteur sequence types (STs), including 2 novel STs first reported by this study (ST1562^Pas^ and ST1563^Pas^). The majority of isolates (77%) belonged to ST2^Pas^ or CC2, the dominant antibiotic-resistant lineage that has circulated in the US and Europe (5). Three sub-lineages within CC2 with varying degrees of heterogeneity were apparent (Figure 1, Table S1). CC2 sub-lineage A (CC2A) comprised multiple Oxford STs, including ST208^Ox^, ST218^Ox^ and ST417^Ox^ (Tables S1, S2). CC2 sub-lineages B (CC2B) and C (CC2C) corresponded to ST451^Ox^ and its single locus variants (SLVs), and ST281^Ox^ and its SLVs, respectively (Table S2).

**Figure 1.**
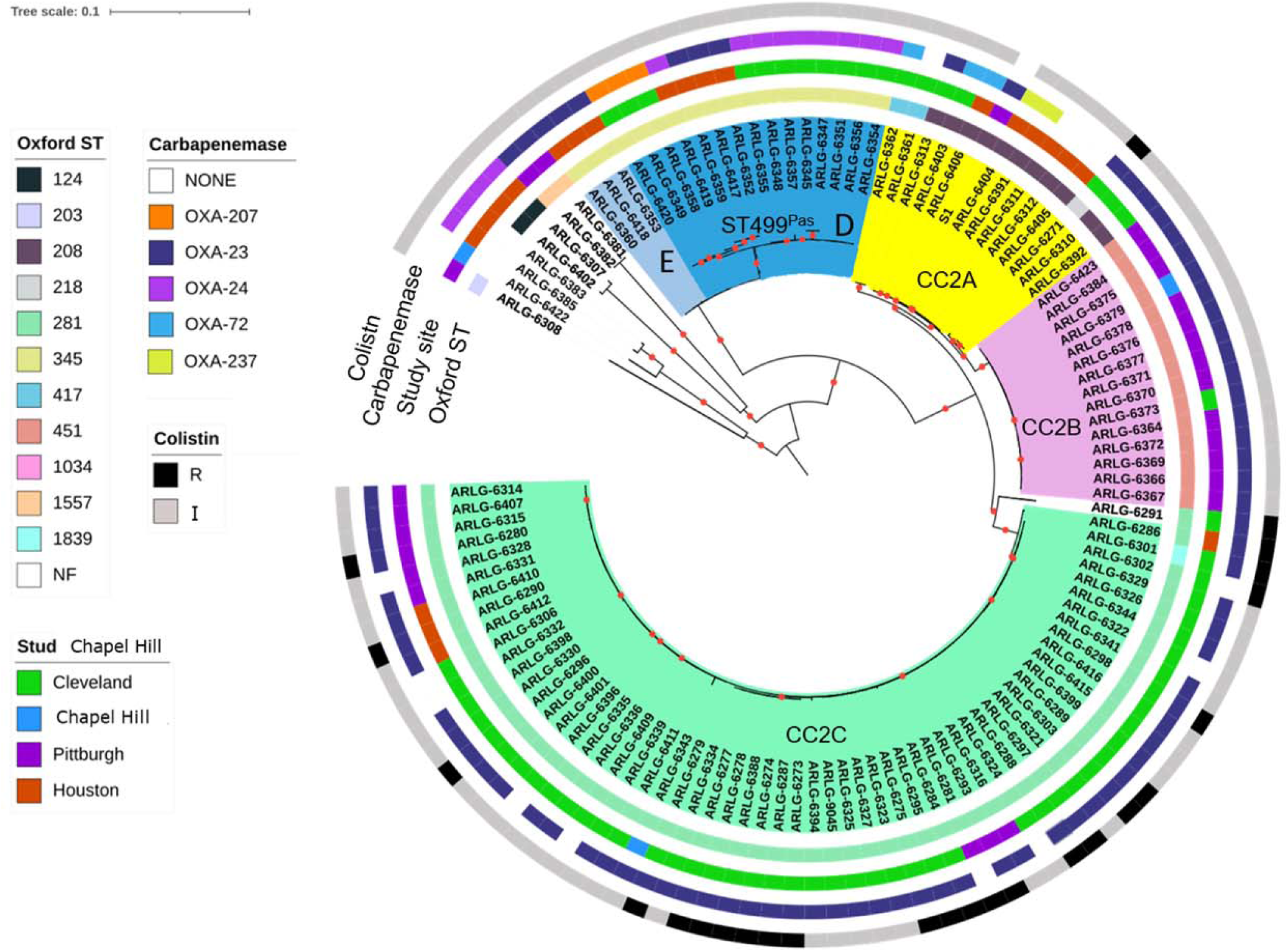
Core genome phylogeny of 115 CR*Ab* isolates from four medical centers in the US. The first isolate sampled from each patient was included, and the mid-point rooted phylogeny was constructed from single nucleotide polymorphisms (SNPs) detected in the core genome of all isolates (2.6 Mb core genome length), using RAxML. The phylogeny is annotated based on Oxford ST and study center of isolation. Branches are shaded by lineages and sub-lineages described in the text. Nodes supported by bootstrap values of 100 are marked with red dots. NF, Not found; R, resistant; I, intermediate. An interactive version of this figure is available online at http://arlg.med.unc.edu/crackle/.

Of the remaining non-CC2 isolates, most belonged to ST499^Pas^ (16%), which has been infrequently observed in the past. The non-CC2 ST499^Pas^ lineage could be further separated into two sub-lineages, D and E (containing 17 and 3 isolates respectively), with both sub-lineages corresponding to the same Oxford ST (ST345^Ox^). The remaining isolates belonged to 7 additional STs and contained one or two isolates each.

The sub-lineage that was previously widely distributed in the US, corresponding to ST208^Ox^ and here called sub-lineage CC2A, comprised a minority of the isolates in our study, and was found only in Cleveland and Houston (Table 2). Isolates belonging to sub-lineage CC2B (ST451^Ox^ and its SLVs) were predominantly found in Pittsburgh, but were also detected in Cleveland and Chapel Hill. Sub-lineage CC2C (ST281^Ox^ and SLVs) was found at all four study centers and was the dominant lineage in Cleveland and Pittsburgh. ST499^Pas^ lineage D isolates were identified in Cleveland and Houston, while lineage E isolates were only found in Houston (Table 2). At the level of individual hospitals, some clinical sites had a single dominant lineage, while others had several dominant lineages. The distributions of CR*Ab* sub-lineages from different body sites were similar to one other, with the exception that no CC2A isolates were found in blood and no CC2B isolates were found in wound cultures (Table 3).

**Table 2.** Geographic distribution of CR*Ab* isolates by study site.

| **Sub-lineages** | **Total (n=115)** | **Cleveland (n=66)** | **Pittsburgh (n=25)** | **Houston (n=21)** | **Chapel Hill (n=3)** |
| --- | --- | --- | --- | --- | --- |
| **CC2A** | 13 (11%) | 7 (11%) | None | 6 (29%) | None |
| **CC2B** | 15 (13%) | 1 (2%) | 13 (52%) | None | 1 (33%) |
| **CC2C** | 60 (52%) | 46 (70%) | 9 (36%) | 4 (19%) | 1 (33%) |
| **ST499<sup>Pas</sup>** | 18 (16%) | 11 (17%) | None | 7 (33%) | None |
| **Other ST** | 9 (8%) | 1 (2%) | 3 (12%) | 4 (19%) | 1 (33%) |

**Table 3.**
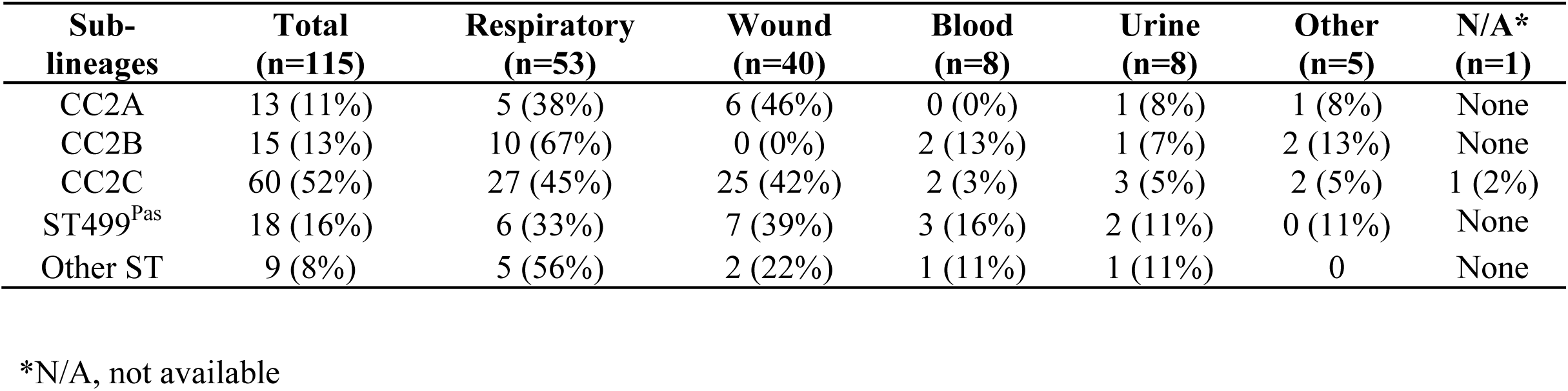
Culture source distribution of CR*Ab* isolates.

### Antibiotic susceptibility of CR*Ab* isolates

We performed MIC testing on the 115 unique patient isolates for agents that possess activity against *A. baumannii*. Of the 115 isolates tested, 36% were resistant to amikacin and 57% were resistant to gentamicin (Figure 2, Table S3). Rates of tigecycline and minocycline resistance were low at 2% and 4%, respectively, whereas 37% of isolates were resistant to cefepime, and 79% were resistant to ceftazidime. Seven isolates (6%) were resistant to cefiderocol using CLSI criteria (MIC, ≥16). Only one additional isolate met resistance criteria when we applied FDA breakpoints (I= 2, R≥4). However, ten isolates (10%) were now considered to have intermediate susceptibility with MIC of 2. Finally, 22% of isolates in our study were resistant to colistin, the last resort antibiotic for treating CR*Ab* infections. Due to the recent change of colistin breakpoints by CLSI, the remaining isolates were intermediate to colistin, eliminating susceptible category and moving susceptible to intermediate. When we assessed antibiotic susceptibility rates by bacterial sub-lineage, a few notable differences emerged. Sub-lineage CC2B had the highest rates of aminoglycoside and cefepime non-susceptibility rates. ST499^Pas^ (D and E) isolates, on the other hand, had comparatively low rates of non-susceptibility to ceftazidime and cefepime. All except two colistin-resistant isolates belonged to sub-lineage CC2C, resulting in 38% resistance within this sub-lineage. Study sites differed somewhat in the proportion of colistin-resistant CC2C isolates: 38% in Cleveland, 33% in Pittsburgh, and 50% in Houston. Most cefiderocol non-susceptible isolates belonged to sub-lineage CC2C as well, and one isolate was resistant to both cefiderocol and colistin. Cefiderocol non-susceptibility within CC2C lineage increased from 6% to 23% when FDA breakpoints were applied.

**Figure 2.**
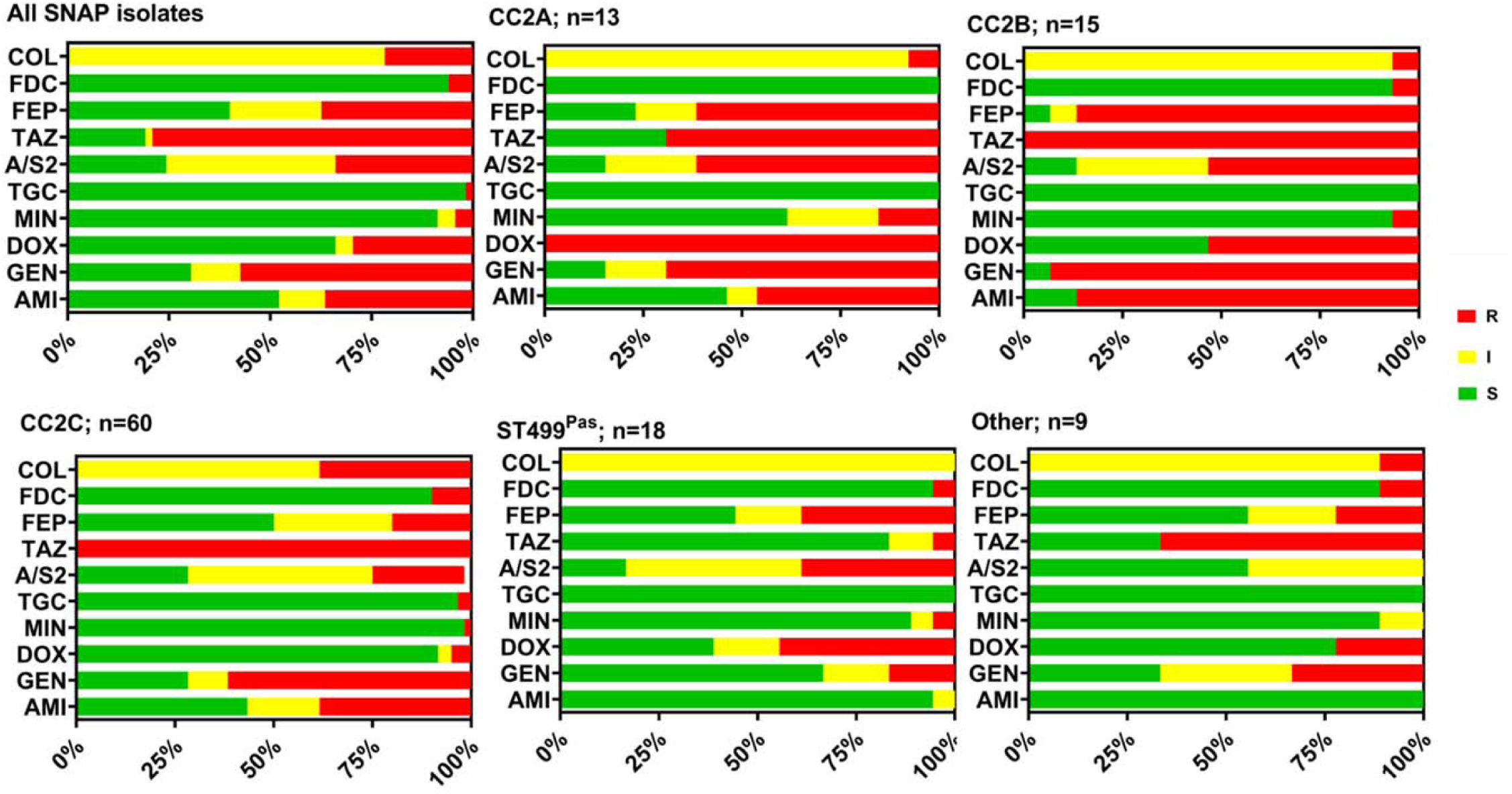
Antimicrobial susceptibility profiles of CR*Ab* isolates. Antimicrobial susceptibilities of 115 initial patient isolates were determined with Sensititre plates or broth microdilution. COL, colistin; FDC, cefiderocol; FEP, cefepime; TAZ, ceftazidime; A/S2, ampicillin-sulbactam; TGC, tigecycline; MIN, minocycline; DOX, doxycycline; GEN, gentamicin; AMI, amikacin. S, susceptible; I, intermediate; R, resistant. Susceptibilities were assigned according to the Clinical and Laboratory Standards Institute (CLSI) guidelines.

### Plasmids and resistance islands

One of the reasons for the success of CR*Ab* as a nosocomial pathogen is its ability to acquire drug resistance genes through horizontal transfer. In addition to the ability to acquire plasmids, CR*Ab* isolates are also known to possess composite transposons and integrons containing resistance genes at chromosomal locations, referred to as resistance islands (RIs) (14, 15).

To determine plasmid diversity within clinical CR*Ab* isolates, six unique plasmids were resolved from available high-quality, closed genomes from isolates belonging to CC2A (S1), CC2B (ARLG-6376), CC2C (ARLG-6295, ARLG-6344), ST499^Pas^ D (ARLG-6420) and ST499^Pas^ E (ARLG-6418). Six unique plasmids were resolved (Table S4). Plasmids varied in size from 11 kb to 167 kb and belonged to five different homology groups based on *rep* gene sequences. Three of the plasmids carried OXA-type carbapenemase genes (*bla*_OXA-23_ in pARLG6295_2 and pARLG6344_2; *bla*_OXA-207_ in pARLG6420_2). pARLG_6295_2 additionally encoded the *aphA6* gene conferring amikacin resistance. The remaining two plasmids did not possess known antimicrobial resistance genes.

Next, we evaluated the presence of identified plasmids among all initial CR*Ab* isolates in our study (Figure 3). Overall, CC2 isolates carried significantly more plasmids than non-CC2 isolates (*p* <0.0001, Mann-Whitney test). Within CC2, different lineages had differences in plasmid content, with most CC2B isolates containing pARLG6344_3, while CC2C tended to harbor the *bla*_OXA-23_-carrying plasmids pARLG6344_2 and pARLG6295_4. CC2C isolates that did not contain pARLG-6344_2 had either the *bla*_OXA-23_ and *aphA6*-encoding pARLG6295_2 plasmid, or no detectable carbapenemase gene-carrying plasmids. The majority of ST499^Pas^ isolates lacked plasmids, with the exception of 3 isolates carrying *bla*_OXA-207_ on pARLG6420_2. To account for additional plasmids, we examined sequences for the presence of plasmid *rep* genes (14). Overall, plasmid *rep* genes belonging to eight groups were identified among all initial CR*Ab* isolates. Plasmid *rep* gene content was also higher in CC2 versus other lineages (*p* <0.0001, Mann-Whitney test).

**Figure 3.**
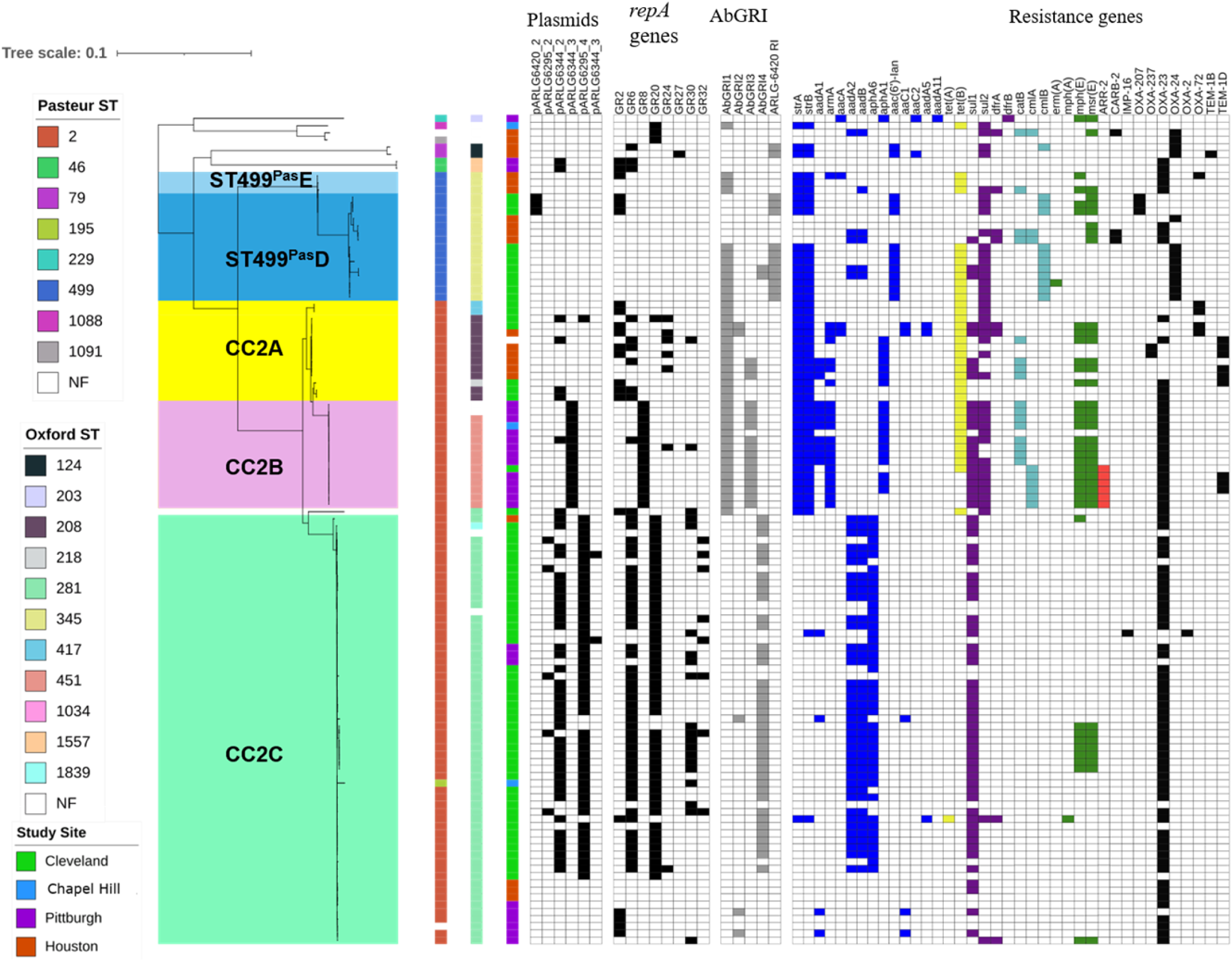
CR*Ab* phylogenetic tree from Figure 1 shown with plasmid content, genomic resistance islands (AbGRI), and antibiotic resistance genes. Plasmids were identified by long-read sequencing and *repA* gene sequence presence. Presence of genetic elements is notated with color boxes; absence is shown with white boxes. Plasmid and β-lactamase gene presence is shown with black boxes. AbGRI presence is shown with grey boxes. Resistance gene presence is shown with colored boxes: blue, aminoglycoside resistance; yellow, tetracycline resistance; purple, sulfonamide resistance; turquoise, chloramphenicol resistance; green, macrolide resistance; red, rifampin resistance.

We then surveyed the isolates for the presence of previously described RIs, including AbGRI1, AbGRI2, AbGRI3, and AbGRI4 (Figure 3) (16). Most CC2A and CC2B isolates, along with some ST499^Pas^ isolates, possessed an AbGRI1-like island, which typically carries *strA-strB* (streptomycin resistance), *tetA*(B) (tetracycline resistance), and *bla*_OXA-23_ genes. Most CC2B and several CC2A isolates belonging to ST2^Pas^/ST208^Ox^ also contained an AbGRI3-like RI carrying *aacA4* (gentamicin/tobramycin resistance), *catB8* (chloramphenicol resistance), *aadA1* (streptomycin resistance), and *armA* (gentamicin, kanamycin, amikacin, tobramycin, and plazomicin resistance). CC2C isolates almost exclusively contained the recently described AbGRI4 island containing *aadB* (tobramycin resistance), *aadA2* (streptomycin and spectinomycin resistance), and *sul1* (sulfonamide resistance) genes. A small group of CC2C isolates lacked AbGRI4 and contained either an AbGRI2-like RI, which typically carried *aacC1* (gentamicin resistance), *aadA1* (streptomycin resistance), and *sul1* (sulfonamide resistance), or no RIs at all.

Additionally, we identified a RI that was exclusively present in ST499^Pas^ and ST79^Pas^ isolates within our dataset. This RI (ARLG-6420 RI) was 19.5-kb long and was integrated at the *tRNA-Ser* site. It possessed 99.8% sequence identity with Tn*6250*, which was previously described in strain LAC-4, an ST10^Pas^ CR*Ab* that was associated with an outbreak in Los Angeles County, CA (Figure 4) (17, 18). Overall, these findings suggest that both plasmids and resistance islands are abundant among CR*Ab* isolates, and they encode clinically relevant antimicrobial resistance genes that likely contribute to the persistence of CR*Ab* in clinical settings.

**Figure 4.**
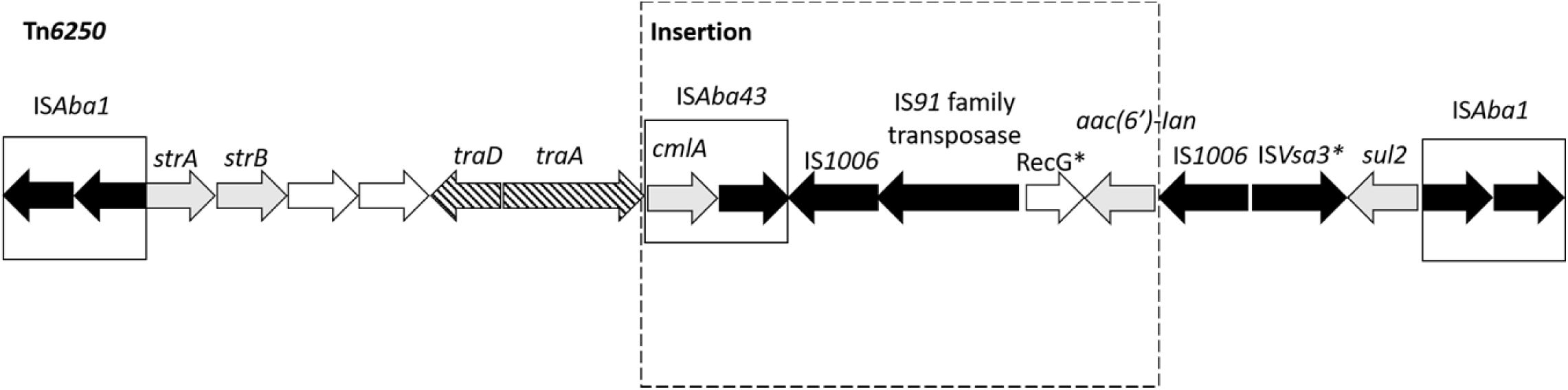
Tn*6250*-like resistance island identified in ST499^Pas^ and ST79^Pas^ isolates. Nomenclature and labeling match the original publication of LAC-4 genome for comparison. Direction of transcription of genes is shown by arrows. Dark arrows denote transposase genes. Gray arrows denote antimicrobial resistance gens. Striped arrows and open arrows denote genes involved in conjugation and genes with unknown function, respectively.

### Carbapenem resistance mechanisms

We catalogued all genomes for carbapenemase genes that would explain their CR*Ab* phenotype. The most frequent acquired carbapenemase gene detected was *bla*_OXA-23_, which was present in 69% of isolates (Figure 3). Other acquired *bla*_OXA_ carbapenemase genes were identified less frequently and included *bla*_OXA-24/40_; *bla*_OXA-72_ and *bla*_OXA-207_ (encoding single amino acid variants of OXA-24/40); and *bla*_OXA-237_ (encoding a recently characterized OXA-235-like carbapenemase) (19-21). Fourteen isolates belonging to different sub-lineages, primarily CC2A and CC2C, did not encode known acquired carbapenemase genes, despite being resistant to carbapenems. Of these 14 isolates, 8 isolates possessed an insertion of IS*Aba1*, an insertion sequence carrying strong promoter activity upstream of the intrinsic carbapenemase gene *bla*_OXA-82_, whose product shows weak carbapenemase activity at baseline expression. The same IS*Aba1* insertion upstream of *bla*_OXA-82_ was previously reported to result in carbapenem resistance in *A. baumannii* (22). Another 4 isolates possessed IS*Aba1* insertions upstream of other chromosomal β-lactamase/carbapenemase genes, including *bla*_OXA-172_, *bla*_OXA-113_ and *bla*_OXA-916_.

### Colistin resistance in CC2C

Given the surprisingly high rate of colistin resistance in sub-lineage CC2C isolates, we explored possible mechanisms of colistin resistance within this sub-lineage. None of the isolates contained *mcr* gene family sequences encoding acquired colistin resistance determinants (23). Additionally, we examined the sequence of the *pmrCAB* operon, which is responsible for lipopolysaccharide (LPS) modifications leading to colistin resistance, as well as the *lpxA, lpxC*, and *lpxD* genes, which are involved in LPS synthesis and whose disruption can result in colistin resistance (24, 25). We found that *pmrA, lpxA*, and *lpxC* had identical nucleotide sequences among both colistin-intermediate and colistin-resistant isolates. Non-synonymous *pmrB* mutations were present in 30% of colistin isolates (L9P, I25F, M145K, F155V, E185K, F387Y, and N353S). We also identified a D95E substitution in *lpxD* in one isolate. The contribution of *pmrC* and *eptA* sequence variation could not be evaluated. For 65% of the isolates, we could not identify a mechanism of colistin resistance based on the analysis of candidate resistance-associated genes.

### Frequency of recombination events

An important question in bacterial population genetics is the extent to which recombination contributes to or constrains lineage diversity. We used ClonalFrameML to analyze all 150 available CR*Ab* genomes (Figure 5A-B) and discovered an overall recombination rate of 64 recombination events for every 100 point mutations. Within CC2 and ST499^Pas^, the rates were 43 and 49 per 100 point mutations, respectively. Major recombination hotspots within CC2 occurred in probable prophage regions and in the capsular polysaccharide locus (Figure 5A). Several other long putative recombination events distinguished different sub-lineages and Oxford STs within CC2 and affected predicted capsular polysaccharide loci and surrounding genes, including *gpi*, which is one of the genes involved in defining Oxford ST (Figure 5A) (26). Similarly, within ST499^Pas^, larger recombination hotspots occurred in predicted prophage regions, as well as in other putative mobile genomic elements (MGEs), including the Tn*6250*-like genomic island described above. However, recombination events were spread out throughout the genome, largely sparing the CPS locus in this clade (Figure 5B). Finally, removal of recombinant SNPs decreased the number of SNPs in the core genomes and merged the CC2A and CC2B sub-lineages and the ST499^Pas^ D and ST499^Pas^ E lineages (Supplementary Tables 6 and 7). These data demonstrate high rates of recombination within CR*Ab* populations that led to differentiation of CR*Ab* sub-lineages both within CC2 and ST499^Pas^.

**Figure 5.**
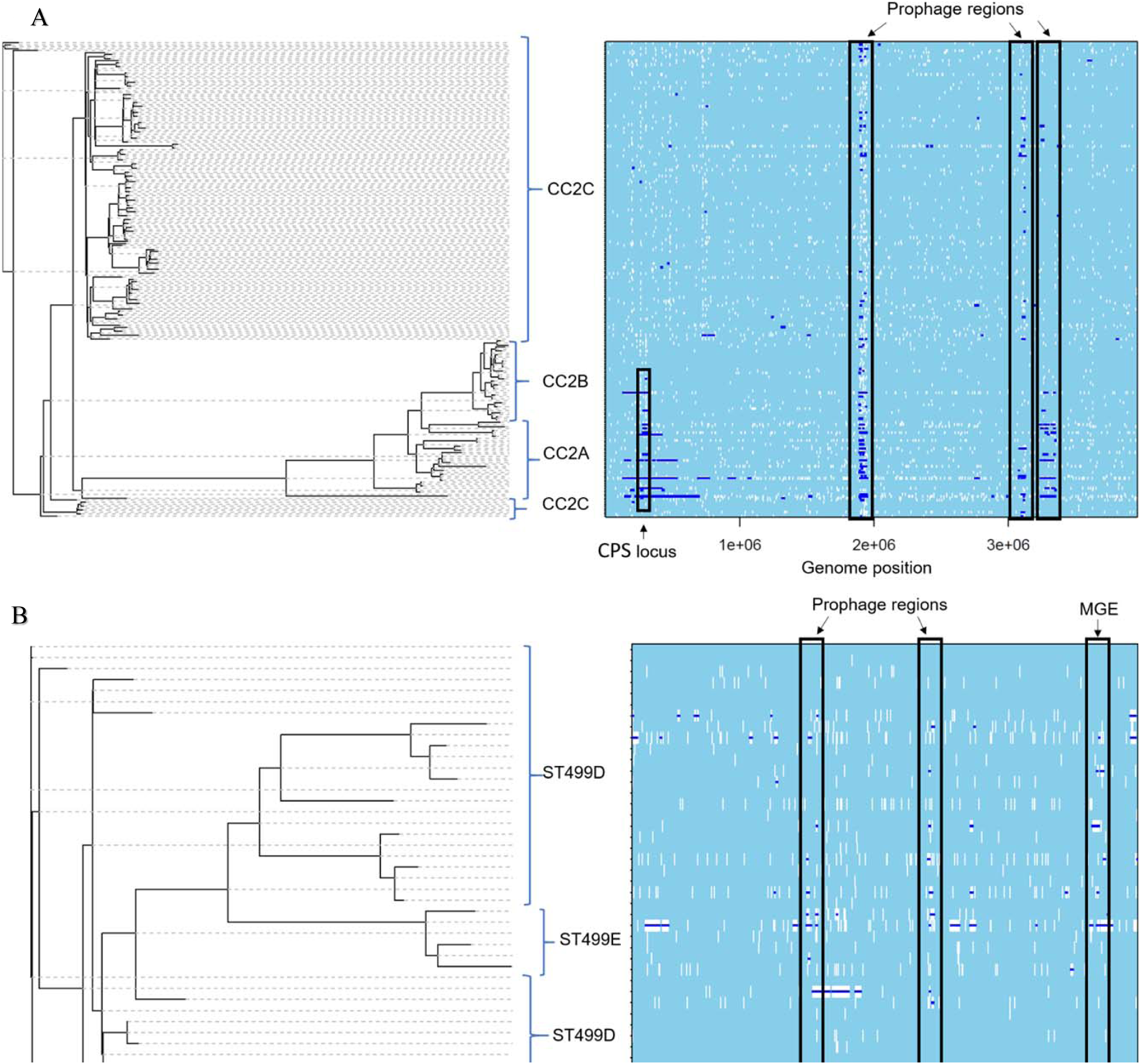
ClonalFrameML analysis of recombination in CR*Ab* lineages CC2 (A) and ST499^Pas^ (B). White vertical bars represent nucleotide substitutions along each branch of the phylogenetic tree. Dark blue horizontal bars indicate putative recombination events. Major CC2 lineages are identified by blue brackets. Recombination hotspots and are identified with black rectangles. The S1 genome was used as a reference for the CC2 lineage, while the ARLG-6345 closed genome was used for the ST499^Pas^ lineage. CPS, capsular polysachride; MGE, mobile genetic element.

### Intra- and inter-patient genetic diversity

We next assessed the genetic diversity of the CR*Ab* isolates within and between patients. Of the 24 patients with more than one isolate from different culture dates, 22 yielded CR*Ab* isolates belonging to the same Oxford ST. The two remaining patients had CR*Ab* isolates belonging to two distinct Oxford STs, suggesting that most patients with multiple isolates were colonized or infected with the same bacterial strain over time, rather than multiple genetically unrelated strains.

Overall, isolates belonging to the same sub-lineage derived from the same patient had a median pairwise SNP distance of 5 (range, 0-31) (Figure 6, Table S5). Within the same-patient group, sub-lineage CC2A, CC2B, and ST499^Pas^D isolates collected from the same patients were very closely related (range, 0-9 SNPs), while sub-lineage CC2C isolates tended to have more SNPs in pairwise comparisons (range, 1-31 SNPs). Higher pairwise SNP differences were observed among isolates from the same hospital, same study site and different study site isolates for all sub-lineages when compared to same patient isolates. Based on these data, we conclude that isolates belonging to CC2A, CC2B, and ST499^Pas^ that are 5-10 SNPs apart, and CC2C isolates that are up to approximately 30 SNPs apart, likely belong to the same strain and SNP differences this small between isolates from different patients may indicate recent transmission, while SNP differences of more than 100 likely indicate infection by unrelated strains.

**Figure 6.**
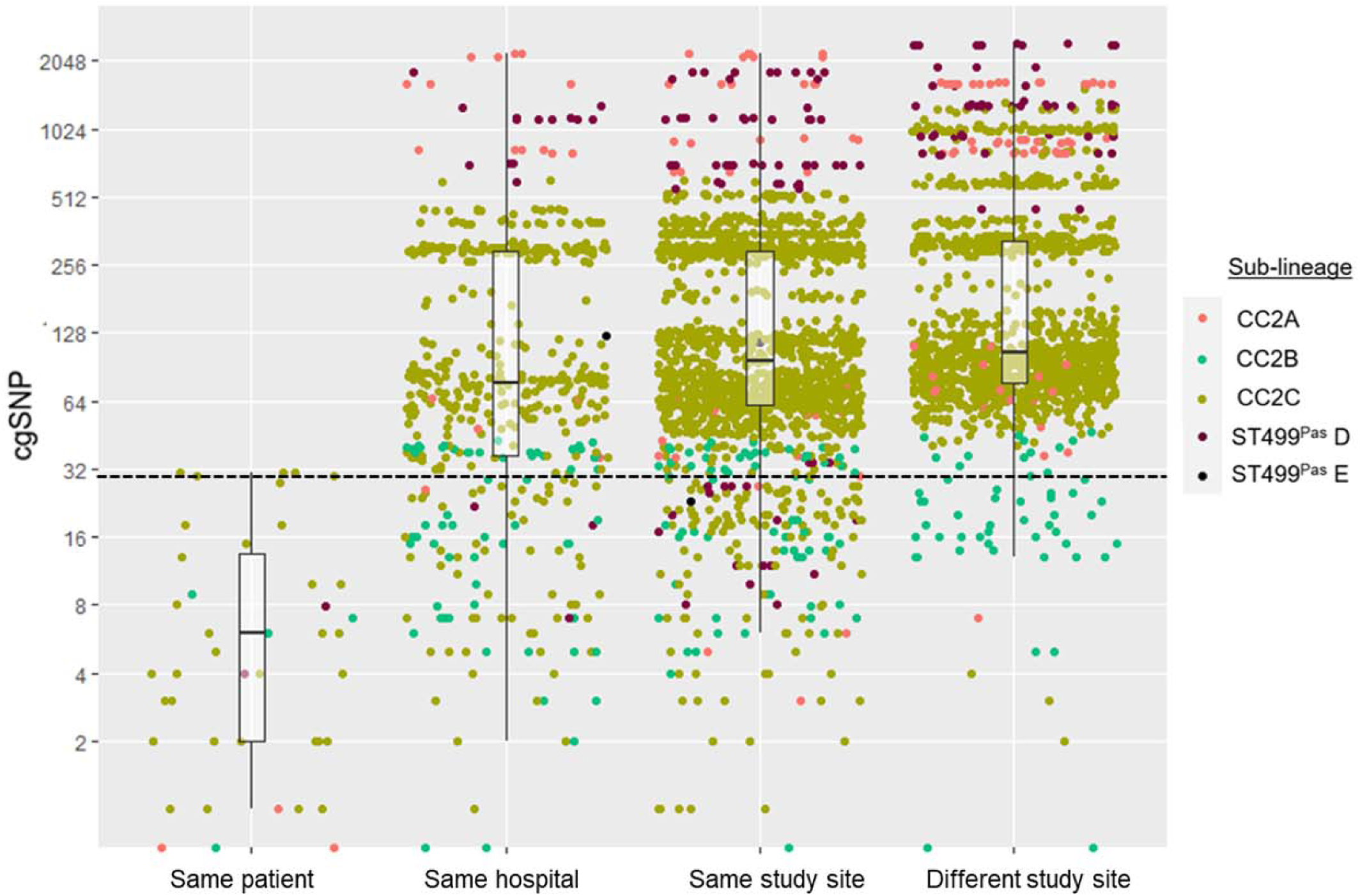
Pairwise core genome SNP distance comparisons between CR*Ab* isolates of the same sub-lineage. A total of 150 isolates from 120 patients were included, and comparisons are color-coded by sub-lineage. Box plots indicate median (horizontal line), interquartile range (box edges), and 1.5 x interquartile range (whiskers) for each group. The dashed horizontal line marks a SNP distance of 31, which corresponds to the maximum pairwise SNP difference observed between isolates sampled from the same patient.

### ST499^Pas^ as an emergent CR*Ab* lineage in the US

ST499^Pas^ was the most common non-CC2 lineage in this study, comprising 16% of all isolates. Since this lineage was not previously known to be a dominant CR*Ab* lineage in the US, we reviewed our findings in the context of ST499^Pas^ isolates genomes deposited in the NCBI database from a variety of locations in the US and from Tanzania (Figure S1). A group of 11 closely related isolates represented an outbreak of CR*Ab* in a Chicago-area hospital between 2009 and 2012 (27). Another 14 isolates, most of which were carbapenem-resistant, were collected in Maryland in 2011 and 2012 (28). The rest of the isolates in the database were reported from Kentucky, Ohio, and Tanzania. Phylogenetic analysis of all ST499^Pas^ genomes from SNAP and NCBI showed a diverse population, with a median pairwise SNP distance of 1,386 (range, 0-7,803) over the 2.3-Mb core genome. The majority of the ST499^Pas^ genomes in the NCBI database were identified as carbapenem-resistant in respective reports, however we were not able to identify previously known carbapenemase genes in 46% (16/35) of isolates. In contrast, all SNAP ST499^Pas^ isolates possessed an acquired carbapenemase gene. Among the combined SNAP and NCBI ST499^Pas^ genomes, *bla*_OXA-24/40_ was the most common acquired carbapenemase gene, followed by *bla*_OXA-23_ and *bla*_OXA-72_ (27, 29, 30). ARLG6420 -RI identified in our cohort, was only found in two unrelated NCBI deposited genomes.

## Discussion

CR*Ab* pose a significant problem worldwide due to their high frequency of multidrug resistance and limited options for effective treatment. In 2019, the CDC Antimicrobial Resistance Threats Report listed CR*Ab* as an urgent public health threat due to limited treatment options and also pointed to their potential to spread carbapenemase genes to non-*Acinetobacter* healthcare-associated pathogens (4). Here, we described the contemporary clinical and genome epidemiology of 120 patients and associated bacterial isolates registered at four major medical centers in the US.

Patients infected or colonized with CR*Ab* were older and were admitted from healthcare settings, such as long-term care facilities. Majority had comorbid conditions, and almost half of the patients were critically ill at the time of initial presentation. Majority of CR*Ab* were isolated from respiratory and wound sources.

Most studies of clinical outcomes of CR*Ab* infection have been derived from observational or retrospective studies, often from single hospital systems. Mortality estimates associated with CR*Ab* infection in the past have been highly variable, ranging between 16% and to 76% (31). In our study, all cause 30- and 90-day mortality rates were 24% and 27%, respectively. In patients determined to have infection, both 30- and 90-day mortality was 26%. These findings underscore the idea that CR*Ab* pose a threat to the most vulnerable patients and contribute to the high morbidity and mortality. Viewed another way, CR*Ab* appear to colonize and infect patients who are at high risk for poor outcomes.

CC2 was the most prevalent CR*Ab* lineage in our study, followed by ST499^Pas^. The two dominant lineages differed in plasmid content, with CC2 isolates having generally more plasmids than non-CC2 isolates, including ST499^Pas^. Additionally, different genomic regions were affected by recombination in CC2 and ST499^Pas^. Most notably, the CPS locus was a hot spot for recombination in CC2 isolates, indicating possible selection for diversification of this trait but also confounding strain assignments based on the Oxford ST scheme. Once these recombination events are accounted for, sub-lineages within CC2 and ST499^Pas^ were no longer distinguishable, demonstrating the role of recombination events in their ongoing diversification. The sub-lineages within CC2 and ST499^Pas^ differed in their geographic distribution, antimicrobial susceptibilities, as well as plasmid and RI content. Overall, these data demonstrate that recombination as well as plasmid and RI content play an important role in the emergence and differentiation of CR*Ab* clonal lineages and their acquisition of antimicrobial resistance determinants (32).

When compared to prior surveillance studies, we saw a shift in the dominant CR*Ab* sub-lineages within CC2. A previous molecular epidemiologic study of CR*Ab* in the US from 2008-2009 showed predominance of ST2^Pas^/122^Ox^ and ST2^Pas^/208^Ox^ lineages among CR*Ab* isolates at six health systems across the US (6). In the present study, we did not identify any ST2^Pas^/122^Ox^ isolates and relatively few ST2^Pas^/208^Ox^ isolates. Instead, sub-lineage CC2C (ST2^Pas^/281^Ox^) emerged as one of the dominant lineages at three of four participating sites. This is consistent with a recent report by Adams et al. describing replacement of ST208^Ox^ by ST281^Ox^ in two Cleveland health systems (16). Similarly, ST281^Ox^ was found to be the dominant sub-lineage at a Maryland hospital in 2011-2012 (28). On the other hand, the emergence of ST499^Pas^ as another dominant lineage was unexpected. While ST499^Pas^ isolates have been reported over the years from different US cities, it has not been considered as an emerging lineage taking a foothold in hospital systems across the country, as was identified in this study. On balance, each hospital system in our study appeared to have its own unique CR*Ab* population. The CR*Ab* populations within individual hospitals were similar to those in the health systems they belonged to. This is not surprising and can be explained by movement of patients and healthcare workers between hospitals and long-term care facilities within the same geographical areas. Some sub-lineages were widely distributed, like sub-lineage CC2C (ST2^Pas^/ST281^Ox^), while others were more localized, like sub-lineage CC2B (ST2^Pas^/ST451^Ox^), which was found primarily in Pittsburgh. Generally, in study sites with many CR*Ab* cases, few clonal sub-lineages dominated, suggesting endemicity within hospitals or associated facilities, as CR*Ab* is known to survive for prolonged periods of time on environmental surfaces such as hospital beds and in sinks and other plumbing (33, 34). While our study was not designed to investigate patient-to-patient transmission, core genome SNP analyses showed that the majority of same patient, same sub-lineage isolates fell within 10 SNPs of one another, whereas isolates from different patients at the same site typically had more than 100 SNPs separating them. This finding may be useful in the interpretation of genome sequencing data of CR*Ab* in the context of hospital epidemiology.

The antibiotic susceptibility profiles of the CR*Ab* isolates we collected were distinct compared to prior surveys. While different sub-lineages had distinct profiles, we observed a general decrease in resistance to cephalosporins and aminoglycosides in our cohort compared to a prior survey, as well as an increase in non-susceptibility to ampicillin-sulbactam (6). A concerning finding was the high rate of colistin resistance. The overall colistin resistance rate in our study was 22%, with sub-lineage CC2C (ST281^Ox^) being the main driver with nearly 40% of isolates being resistant. This is in contrast with a recent study reporting a colistin resistance rate of 8.7% among meropenem-non-susceptible *A. baumannii* isolates collected from hospitals in North America in 2014, as well as the colistin resistance rate of 16.6% reported in a recent study from Europe (35, 36). The fact that we uncovered diverse mutations in *pmrB* and *lpxD* among the colistin-resistant isolates, and that resistant isolates were interspersed throughout the sub-lineage CC2C phylogeny, suggests that colistin resistance probably evolved *de novo* in each patient, rather than spread through transmission of a single resistant clone. It is also possible that sub-lineage CC2C (ST281^Ox^) is more adept at maintaining colistin resistance, which may have lead to overrepresentation of colistin-resistant isolates belonging to this sub-lineage. This finding has implications for empiric treatment options, since colistin or polymyxin B is still often the mainstay of therapy, and susceptibility testing is delayed given the complexity of current testing strategies. Nonetheless, these findings highlight the need for development of new testing strategies for rapid determination of colistin susceptibility, along with alternative therapies, to minimize the risk of administering inactive therapy.

One novel therapy that could be useful for the treatment of CR*Ab* is cefiderocol, a siderophore cephalosporin approved in 2019 after the conduct of this study. Approximately 6% of the CR*Ab* isolates were found to be resistant to cefiderocol using CLSI breakpoints, which is comparable with previously reported surveillance data (35). When we used FDA established breakpoints, the resistance rate largely did not change, however, additional ten isolates were considered to be intermediate to cefiderocol. This finding, in conjunction with recent evidence of higher all-cause mortality in patients who were infected with CR*Ab* and treated with cefiderocol in the CREDIBLE-CR trial (which compared cefiderocol to the best available therapy for carbapenem-resistant organisms), suggests that the role of cefiderocol in the treatment of invasive CR*Ab* infections remains unclear (37). However, it might still be considered as a rescue therapy in critically ill patients, in which case susceptibility of the isolate should be confirmed (38).

This study had several limitations. We only included patients and isolates from four large hospital systems that were the initial participants of the SNAP cohort. These sites were largely self-selected and might not be representative of the US population as a whole. Additionally, this study was restricted in time to 12 months and was thus unable to examine changes in CR*Ab* populations over time. Furthermore, the sample size was relatively small, which makes it difficult to draw clinically relevant conclusions regarding lineage-specific outcomes. We expect to address many of these limitations in future studies of the full SNAP dataset. Systematic studies of CR*Ab* are of paramount importance for devising strategies to prevent their dissemination and improve clinical outcomes. The full SNAP dataset will provide a robust resource for studying CR*Ab* biology and associated clinical outcomes in a systematic, clinically-informed manner.

Overall, our findings highlight the continued importance of CR*Ab* as a healthcare-associated pathogen causing high morbidity and mortality and with limited treatment options, as well as the importance of real-time surveillance and genomic epidemiology in studying their dissemination and clinical impact.

## Acknowledgements

The research reported in this publication was supported by the National Institute of Allergy and Infectious Diseases the National Institutes of Health under Award Number R21AI135522 and in part under Award Number UM1AI104681. The content is solely the responsibility of the authors and does not necessarily represent the official views of the National Institutes of Health. A.I. was supported through Physician Scientist Incubator Program at the University of Pittsburgh sponsored by the Burrows Wellcome Fund. R.A.B. was supported by the National Institutes of Health under Award Nos. R01AI100560, R01AI063517, and R01AI072219. This study was also supported in part by funds and/or facilities provided by the Cleveland Department of Veterans Affairs, Award Nos. and 1I01BX001974 to R.A.B. from the Biomedical Laboratory Research & Development Service of the VA Office of Research and Development, and the Geriatric Research Education and Clinical Center VISN 10. V.S.C. was supported by a research grant from the National Institutes of Health (U01AI124302) and, in part, by PA CURES Grant #4100085725 with the Pennsylvania Department of Health. D.V.T. was supported by a research grant from the National Institutes of Health (R00EY02822). Y.D. was supported by research grants from the National Institutes of Health (R01AI104895, R21AI151362). SSR reports prior employment by bioMerieux (Aug 2019 – May 2021) and research funding from BD Diagnostics, Hologic, Diasorin, and Lifescale while employed in the department of Laboratory Medicine at Cleveland Clinic. CL was supported by a research grant from the National Institutes of Health T32GM086330. CAA was supported by NIH/NIAID grants R01-AI148342-01, R01AI134637, K24-AI121296 and P01-AI152999-01

We thank Shionogi for provision of materials required for susceptibility testing of cefiderocol.

